# *Ovarify*: a generalizable deep learning method for the automatic characterization of ovarian morphology on pelvic ultrasonography

**DOI:** 10.64898/2026.11.16.761040

**Authors:** Eeshaan Rehani, Bailey Smith, Matthew Brendel, Jeffrey Pea, Adam Evans, Faith Carter, Jonas Malmsten, Sujata Kar, Donna Chizen, Roger Pierson, Steven Spandorfer, Kathleen Hoeger, Marla Lujan, Iman Hajirasouliha

## Abstract

Ovarian morphology on ultrasonography is commonly used in the assessment of female reproductive health, such as in the diagnostic evaluation of polyendocrine metabolic ovarian syndrome (PMOS) and monitoring of response to assisted reproduction. Standard morphological markers quantified on pelvic ultrasound are ovarian volume (OV) and measures of antral follicle count (AFC) such as follicle number per slice or per ovary (FNPS, FNPO). Manual assessment of these features is operator-dependent, which limits the reliability of these measurements, impacting the delivery of consistent care. To address this, we have developed *Ovarify*, the first deep learning workflow to automatically characterize ovarian morphology from pelvic ultrasonography across multiple clinical contexts. *Ovarify* was trained on transvaginal ultrasonographic volumes garnered across three centers and evaluated on a held-out internal test set in addition to three external datasets spanning heterogeneity in image collection protocols, scanner hardware, and populations, across both single frames as well as whole-volumes. *Ovarify* segmented the ovary and follicles with high fidelity, outperforming U-Net and ResNet-34 baselines (p<0.001). Morphological measurements agreed closely with validated image analysis protocols on an internal test set (OV: Pearson *r*=0.892, FNPS: *r*=0.667, FNPO: *r*=0.770); across external datasets, workflow performance was similar. *Ovarify* additionally retained performance when evaluated on unseen images of ovaries undergoing controlled ovarian stimulation (FNPS: Pearson *r*=0.813). *Ovarify*-derived measurements classified polycystic ovarian morphology with an AUPRC of 0.90 (F1=0.93) using FNPO and 0.80 using OV. Ultimately, *Ovarify* provides a reproducible, operator-independent characterization of ovarian morphology directly from routinely-acquired pelvic ultrasonograms, generalizing across scanners, acquisition formats, and clinically-distinct imaging contexts. *Ovarify* automates the assessment of clinically-relevant morphological markers, allowing for the unbiased and consistent evaluation of morphological features.

## Introduction

Ovarian morphology assessed using pelvic ultrasonography is a key indicator of female reproductive health. Ovarian size and antral follicle counts inform the evaluation of endocrine disorders such as polyendocrine metabolic ovary syndrome (PMOS), the assessment of ovarian reserve and fertility, and the monitoring of ovarian response during assisted reproductive technology (ART) ^1–3^. Beyond diagnosis, ovarian morphology has been shown to correlate with cycle irregularity, hirsutism, hyperandrogenism, anti-müllerian hormone levels, and measures of hypothalamic-pituitary-ovarian dysfunction; these associations can reflect reproductive symptomology in settings where biochemical assays are inaccessible or clinical assessment is subjective^4–9^.

Ovarian morphology can be quantified through several metrics, including ovarian volume (OV), and antral follicle count (AFC) reflected by follicle number per two-dimensional slice (FNPS) or follicle number per entire (single) ovary (FNPO) ^10^. These measurements reflect underlying ovarian structure and function, and are incorporated into established diagnostic and monitoring frameworks such as the 2023 International Evidence-based Guideline for diagnosis of PMOS, which defines polycystic ovary morphology (PCOM) as FNPO ≥ 20 follicles (or FNPS ≥ 10 follicles), or OV ≥ 10 cm^3^ ^11^. The metrics are additionally reflected in other clinical contexts: in fertility evaluation, AFC is a marker of ovarian reserve, with counts below 5-7 follicles indicating diminished reserve and a poor expected response to ovarian stimulation^12^. In assisted reproduction, serial measurement of follicle number and size guides cycle management, with final oocyte maturation typically triggered once multiple follicles reach a size threshold^13–15^.

Pelvic ultrasonography is the primary modality for evaluating ovarian morphology and is widely accessible and well-suited to point-of-care assessment. Reported diagnostic accuracy for the individual metrics is high, with pooled estimates for FNPO in PMOS ranging from 0.86 to 0.92^2, 11, 16^. In practice, however, the reliability of these measurements is limited by the manual, operator-dependent nature of ultrasound interpretation. The assessment of ovarian features on ultrasonography shows moderate to poor reproducibility; inter-rater agreement for PCOM classification has been estimated at 51%^17, 18^. This has been previously addressed through the development of offline grid-analysis tools that yield more accurate, standardized follicle counts than conventional real-time scanning^19^. However, the specialized training required, and time-intensive nature of offline analysis have limited its clinical adoption, and commercial semi-automated platforms that estimate OV or count and measure follicles have seen slow uptake as their detection routines are tuned for specific clinical scenarios and additionally require manual input^20–22^. Methods that balance accuracy and feasibility are therefore needed for reliable, reproducible characterization of ovarian morphology across clinical scenarios.

Machine learning offers an opportunity to harmonize the assessment of ovarian morphology. Convolutional neural networks (CNNs) are well-suited to extracting the hierarchical features that map onto the structural characterization of the ovary and have been previously applied to ovarian ultrasound for follicle segmentation and counting, biomarker estimation, and even direct classification of PMOS^23–30^. Previously published models, however, face several drawbacks. First, most studies optimize discrimination of diagnostic categories while leaving clinical utility, i.e. interpretability of outputs, unexamined. Second, many models are trained and benchmarked on small, homogeneous datasets with poor label quality, which inflates apparent performance and limits generalizability to routinely acquired clinical scans, including to multiple acquisition formats such as 2D single frames as well as 3D volumes^29, 31–33^. Further, previous models are not validated against external cohorts^20, 24, 25, 27, 33^. An automated platform for ovarian morphology characterization, trained and validated on multiple well-characterized populations utilizing reproducible image-analysis approaches for label generation, would therefore streamline the interpretation of transvaginal ultrasounds.

To address this, we have developed *Ovarify*, a deep learning method that automatically characterizes ovarian morphology from pelvic ultrasonography. The *Ovarify* workflow identifies ovarian content and delineates ovarian structures directly from routinely-acquired transvaginal ultrasounds, and subsequently reproducibly calculates morphological features. *Ovarify* is operator-independent, reducing the user-specific biases that currently constrain the assessment of ovarian morphology.

## Results

### Ovarify

We developed *Ovarify*, a novel deep learning workflow that explainably characterizes ovarian morphology directly from pelvic ultrasonography and outputs segmentation masks for the ovary and antral follicles, in addition to clinically-relevant morphological measurements: ovarian volume (OV), follicle number per single slice (FNPS), and follicle number per ovary (FNPO). *Ovarify* is organized into three sequential stages that are analogous to the manual clinical analysis workflow. First, a frame-level classifier screens each ultrasound and retains only frames containing ovarian content, reducing an entire volumetric acquisition (if taken) to the region of interest without manual frame selection (Figure 1a). Next, the retained frames are passed to an ovarian segmentation stage that delineates the ovarian boundary (Figure 1b). Lastly, a follicle segmentation stage, initialized from the ovarian segmentation model, resolves individual antral follicles within the ovary (Figure 1c).

**Figure 1.**
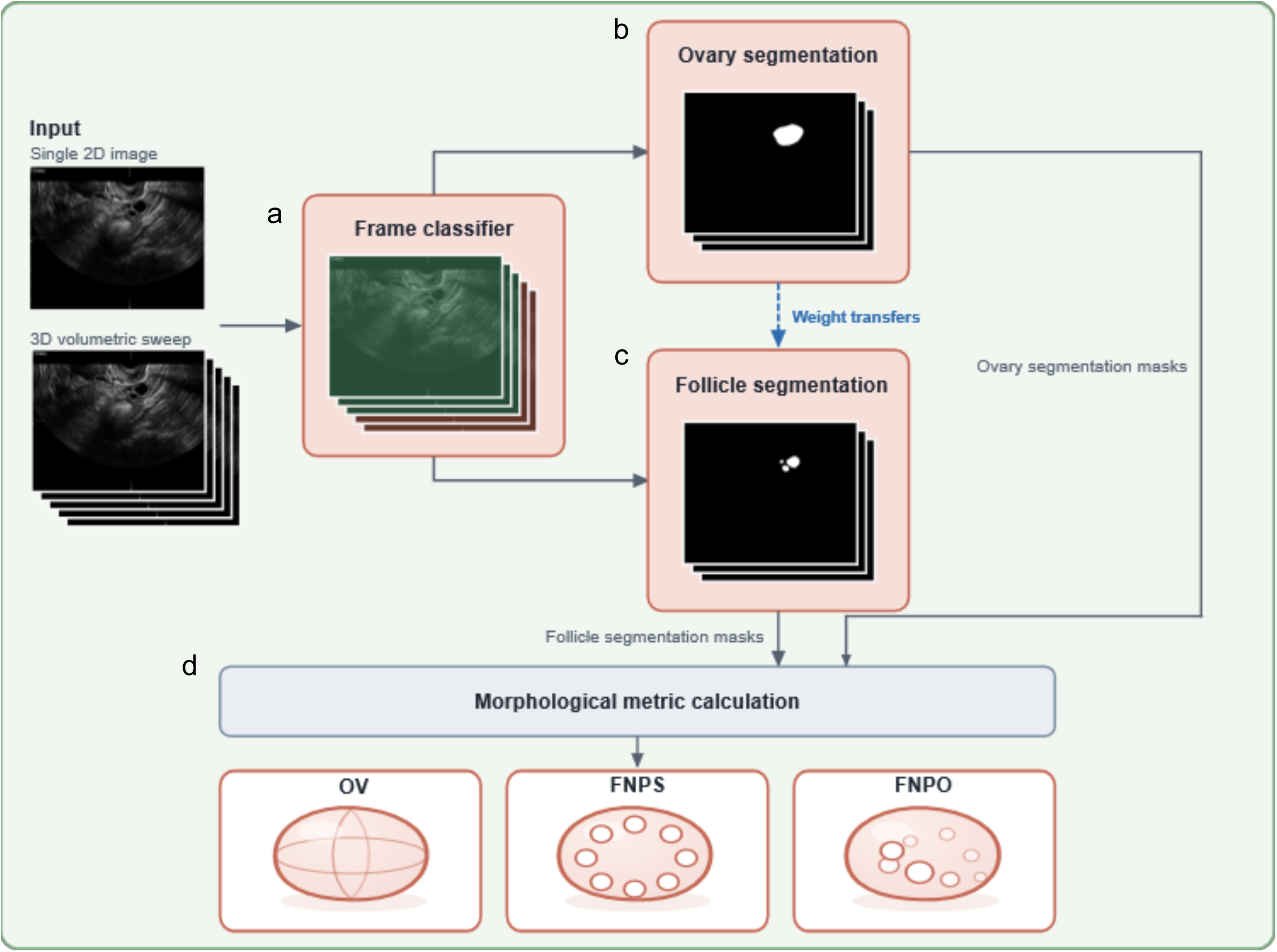
*Ovarify* workflow. (a) *Ovarify* takes as input transvaginal ultrasounds as either static single-frames or volumes, and first classifies frames as either having or not having ovarian content. (b) *Ovarify* segments the ovary from positively-classified frames. (c) *Ovarify* segments antral follicles from positively-classified frames. Training *Ovarify* follicle segmentation employs a transfer learning approach with weights initialized from converged ovarian segmentation. (d) *Ovarify* applies the predicted ovarian and follicle segmentation masks to calculate ovarian volume (OV), follicle number per slice (FNPS), and follicle number per ovary (FNPO).

The segmentation outputs are used to derive clinical measurements through a post-processing step rather than regressed directly, enhancing interpretability of the morphological metrics. Ovarian volume is computed by integrating segmented ovarian area across the acquisition, while follicle counts are derived from the follicle masks in two and three dimensions to yield FNPS and FNPO, respectively (Figure 1d). Because the workflow operates on individual frames, *Ovarify* applies to both single two-dimensional images as well as three-dimensional volumetric sweeps within a single framework. producing per-slice measurements from static frames and volumetric measurements by aggregating across a sweep. This design yields reproducible, operator-independent characterization of ovarian morphology from routinely acquired ultrasonography.

#### Characteristics of datasets used

The primary (Cornell) dataset comprised 986 3D transvaginal ultrasonograms collected from 284 participants across three centers (Cornell University, Weill Cornell Medicine, and University of Rochester Medical Center), all of which were acquired on GE Voluson E or S-Series systems under a standardized image collection protocol, consisting of healthy and polycystic ovaries. The dataset was randomly partitioned at the participant level into a training subset of 240 participants (856 unique scans, comprising the entire *Ovarify* training set) and a held-out test subset of 44 participants (130 scans). The training set comprised images collected from women that had a mean age at ultrasound of 27.9 ± 6.6 years and a mean BMI of 28.7 ± 8.0 kg/m^2^. The held-out test participants had a mean age of 27.9 ± 5.5 years and a mean BMI of 28.9 ± 9.3 kg/m^2^. Both subsets were racially and ethnically diverse; the majority of participants identified as White (64.2% training, 61.4% test) and as not Hispanic or Latino (80.4% training, 72.7% test) (Supplementary table 1, Supplementary table 2). Each stage of the *Ovarify* workflow was trained on annotated frames derived from the training scan set, comprising 75,359 frames for classification, 11,374 frames for ovarian segmentation, and 4,718 frames for follicle segmentation. The held-out test set, reserved for evaluation across all stages, consisted of 10,917 frames for classification, 1,600 frames for ovarian segmentation, and 741 frames for follicle segmentation.

Beyond the primary test dataset, *Ovarify* was additionally evaluated on four external datasets spanning distinct sites, scanner models, and populations. The *External_1* dataset comprised 44 ultrasounds (3,519 frames) from 33 participants with polycystic ovaries; the *External_2* dataset comprised 70 ultrasounds (3,150 frames) from 53 participants with healthy or polycystic ovaries; the *External_3* dataset comprised 54 ultrasounds (7,258 frames) from 33 participants with healthy or polycystic ovaries presenting with fertility concerns; the *External_4* dataset comprised 367 single-frame ultrasounds (367 frames) from patients undergoing controlled ovarian stimulation.

All datasets were annotated by trained annotators using a prescribed image analysis protocol, and during image analysis protocol training, were evaluated for annotation performance. An internal reliability study confirmed that annotators were consistent in their delineation of ovarian and follicular content (Supplementary table 3).

#### Frame classification, ovarian segmentation, and follicle detection

We first evaluated *Ovarify* in its single-frame configuration, in which each frame is processed independently without inter-frame context. The frame-level classifier without three-dimensional context reliably distinguished frames containing ovarian content from those without. Across four folds, the classifier achieved a mean area under the receiver operating characteristic (AUROC) of 0.969 (95% CI, 0.963-0.975), accuracy of 0.910 (95% CI, 0.895-0.925), and recall of 0.887 (95% CI, 0.833-0.941). The classifier significantly outperformed a baseline ResNet-34 across AUROC (baseline, 0.965 [95% CI, 0.962-0.968], p<0.001) and accuracy (baseline, 0.891 [95% CI, 0.870-0.912], p<0.001); the baseline however achieved a higher recall compared to the *Ovarify* classifier (baseline, 0.900 [95% CI, 0.856-0.944], p=0.006) (Figure 2a, Supplementary table 4). On the retained frames, ovarian segmentation achieved a mean Dice score of 0.818 (95% CI, 0.809-0.827), and follicle segmentation achieved a mean Dice score of 0.660 (95% CI, 0.644-0.676). Both segmentation stages statistically significantly outperformed a baseline U-Net trained to convergence with the largest margin observed for follicle segmentation, presumably due to the relative difficulty of the task as compared to ovarian segmentation (ovary segmentation baseline, 0.793 [95% CI, 0.783-0.803], p<0.001; follicle segmentation baseline, 0.506 [95% CI, 0.493-0.519], p<0.001) (Figure 2b, Supplementary table 5).

**Figure 2.**
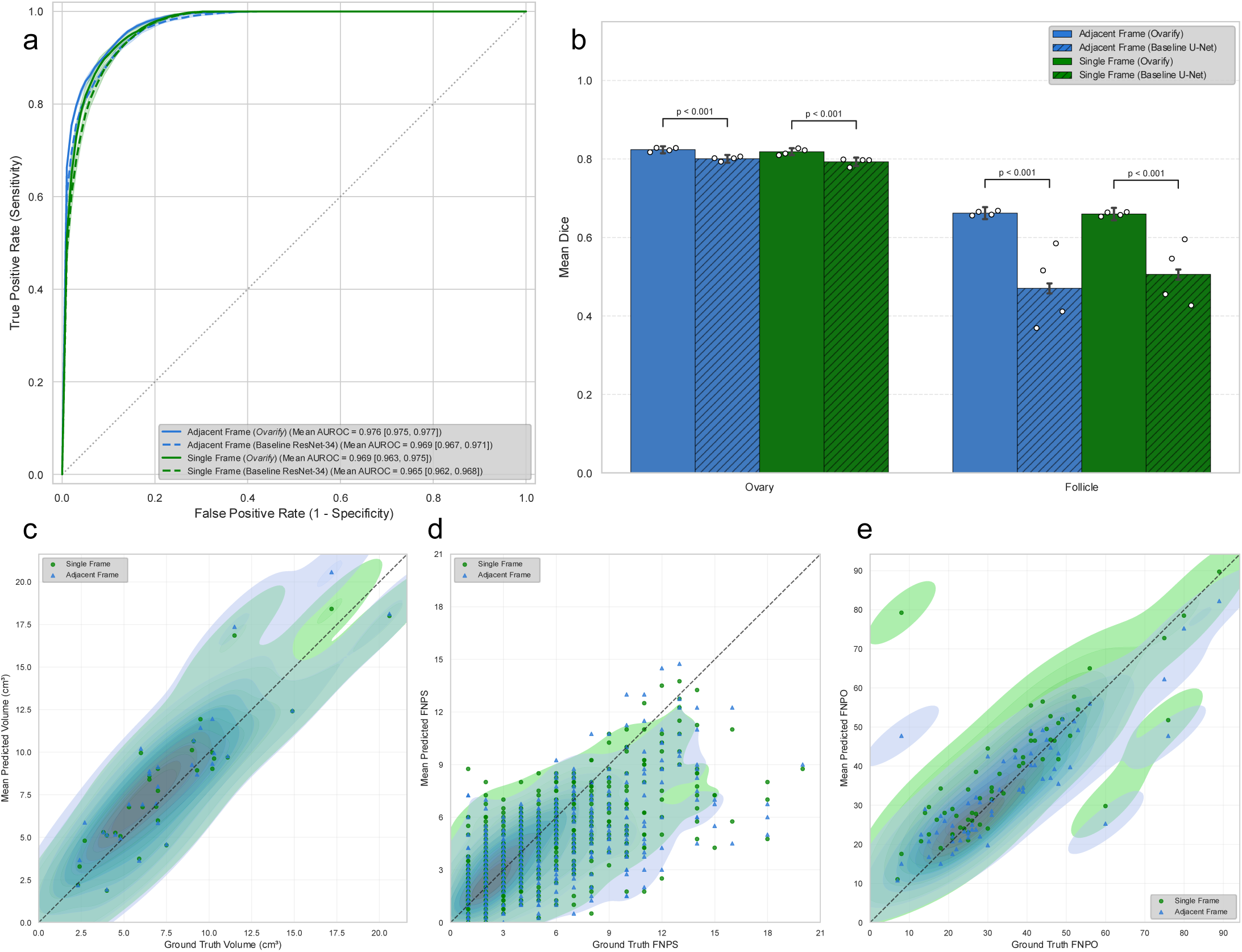
Adjacent-frame and single-frame performances of *Ovarify* on Cornell held-out test set. (a) Frame classification performance of *Ovarify* compared to baseline ResNet-34 classifier. Diagonal dashed line represents random chance predictions. (b) Ovary and follicle segmentation performance of *Ovarify* compared to baseline U-Net segmentation model. Error bars represent 95% confidence intervals across 4-fold cross-validation. Points represent mean Dice score for individual folds. (c) *Ovarify-*derived ovarian volume calculation performance on subset of Cornell held-out test set with available pixel-cm conversion factor and grid-analysis-based ground truth. (d) *Ovarify-*derived FNPS performance on all frames in Cornell held-out test set. (e) *Ovarify*-derived FNPO calculation on fully-labeled subset of Cornell held-out test set. (c-e) Dashed lines represent perfect agreement.

*Ovarify* segmentation outputs translated into accurate morphological measurements. Within the Cornell held-out test set, for all ovaries for which there was a recorded pixel-cm conversion factor as well as ground-truth grid-analysis-based OV (n=31 ovaries), predicted OV agreed with manual measurement at a mean Pearson *r* of 0.892 across folds, with a mean absolute error (MAE) of 1.57 cm^3^. Follicle count across frames (n=741 frames) agreed with a Pearson *r* of 0.667 (MAE 1.62 follicles/slice), and FNPO, evaluated on a fully-annotated held-out subset (n=61 ovaries) agreed with a Pearson *r* of 0.770 (MAE of 6.82 follicles/ovary) (Figure 2c-e).

Considering the 2023 International Evidence-based Guideline for diagnosis of PMOS, we sought to understand whether *Ovarify*-calculated morphological markers aligned with clinical recommendations for the identification of polycystic ovary morphology^11^. In applying model-calculated FNPO, *Ovarify* correctly identified all polycystic ovaries (50 ovaries identified as having ≥20 follicles per ovary), achieving an area under the precision-recall curve (AUPRC) of 0.903 and F1-score of 0.926. Within a margin of 3 counted follicles, *Ovarify* achieved an F1-score of 0.952; excluding ovaries outside of the 3-follicle error threshold (n=10), *Ovarify*-applied FNPO counts achieved an AUPRC of 0.919. Additionally, within a 3-follicle margin, *Ovarify*-applied FNPO counts achieved a balanced accuracy of 0.773 in classifying PCOM. When applying model-calculated OV to classify PCOM, *Ovarify* achieved an AUPRC of 0.799 and an F1-score of 0.533. Within a margin of 1 cm^3^, F1-score increased to 0.933, and AUPRC of PCOM classification increased to 0.963 when considering only ovaries outside of the 1 cm^3^ error threshold (n=24). Balanced accuracy for *Ovarify*-derived OV application to PCOM classification additionally increased from 0.685 to 0.938 when incorporating a 1 cm^3^ volume calculation margin.

We then examined the contribution of ovary-to-follicle weight transfer. We hypothesized that the *Ovarify* follicle segmentation stage trained using weights initialized from the converged ovarian segmentation weights would result in stronger segmentation performance than an equivalent stage trained to convergence with random weight initialization. We found that the incorporation of ovarian segmentation weights into follicle segmentation training significantly improved follicle Dice in the single-frame configuration (random weight initialization, 0.644 [95% CI, 0.628-0.660], p<0.001) (Supplementary figure 1, Supplemental table 6). This is consistent with the anatomical containment of follicles within the ovarian boundary and supports the sequential training philosophy adopted by *Ovarify* (Figure 3).

**Figure 3.**
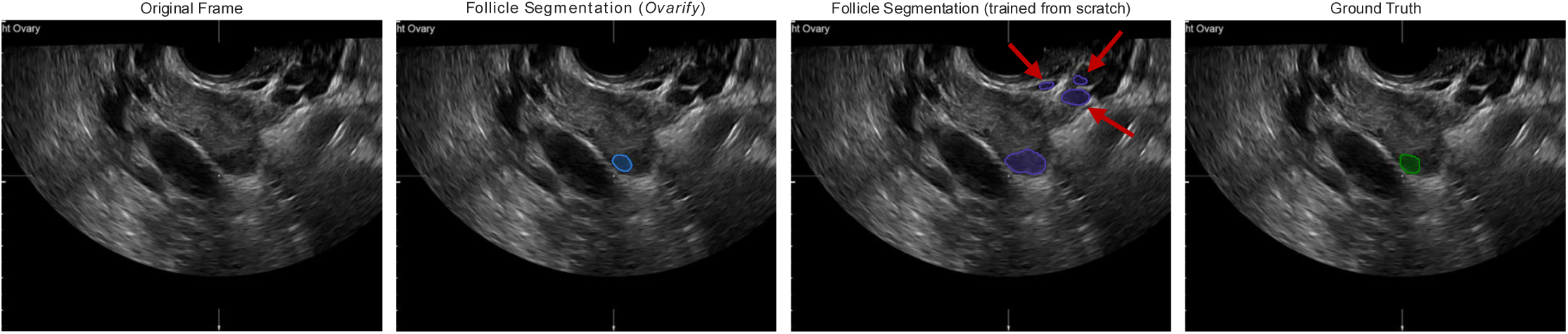
Example follicle segmentation output from *Ovarify*. Weight transfer follicle segmentation training scheme improves *Ovarify* follicle segmentation performance compared to training from scratch by limiting identification of false positive structures (red arrows) outside the ovary. Experienced rater feedback indicates that the pictured frame is considered “difficult”, with an unclear ovarian border.

Finally, we asked whether adding three-dimensional context improved characterization by comparing the single-frame configuration against an adjacent-frame configuration, in which three contiguous frames were stacked depth-wise so that each prediction could draw on its neighboring slices. The two schemes performed comparably on the deep learning tasks. Classification performance and segmentation performances were near-identical (Figure 2a-e, Supplementary table 4, Supplementary table 5), and the downstream morphological measurements showed no significant difference between schemes for OV (MAE 1.57 vs. 1.73 cm^3^, p=0.12), FNPS (MAE 1.62 vs. 1.62 follicles/slice, p=0.94), or FNPO (MAE 6.82 vs. 6.23 follicles/ovary, p=0.73). Classification of PCOM via FNPO achieved an AUPRC of 0.958, F1-score of 0.951, and balanced accuracy of 0.808 within a 3-follicle margin. Classification of PCOM via OV achieved an AUPRC of 0.963, F1-score of 0.875, and balanced accuracy of 0.916 within a 1-cm^3^ margin. These results indicate that although three-dimensional context does not significantly improve model performance in an equivalent workflow, *Ovarify* is notably robust to input scan format, producing similar morphological characterizations whether applied to isolated two-dimensional frames or to frames drawn from a three-dimensional volumetric sweep with adjacent frame context.

#### Validation on external datasets

We evaluated *Ovarify* on three external datasets by retraining the 3D workflow on the entirety of the training (Cornell) dataset. These external datasets span different sites, scanner models, and populations. *External_1* comprised 44 volumes of polycystic ovaries, yielding 3,519 frames collected using a GE Voluson E-Series systems. *External_2* comprised 54 volumes of both healthy and polycystic ovaries, yielding 7,258 frames collected using GE Voluson S-Series scanners. *External_3* comprised 70 manual cineloops of both healthy and polycystic ovaries, yielding 3,150 frames acquired on the Ultrasonix Sonix RP system. Frame classification generalized well across all sites, with an AUROC of 0.965 (External_1), 0.995 (External_2), and 0.988 (External_3) (Figure 4a, Supplementary table 7). Ovarian segmentation was robust, achieving a mean Dice of 0.796 (External_1), 0.817 (External_2), and 0.867 (External_3), while follicle segmentation Dice ranged from 0.653 to 0.690 across sites, consistent with internal performance (Figure 4b, Supplementary table 8). Morphological measurements transferred strongly: predicted OV correlated with manual measurement with a Pearson r of 0.928 (External_1), 0.947 (External_2), and 0.960 (External_3) (Figure 4c-e, Supplementary table 9). Additionally, FNPS correlated with a Pearson r of 0.824 (External_1; MAE 1.465 follicles/slice), 0.803 (External_2; MAE 1.130 follicles/slice), and 0.729 (External_3; MAE 1.563 follicles/slice) (Figure 4f, Supplementary table 10). These results indicate that *Ovarify* generalizes to imaging conditions and populations outside its training distribution.

**Figure 4.**
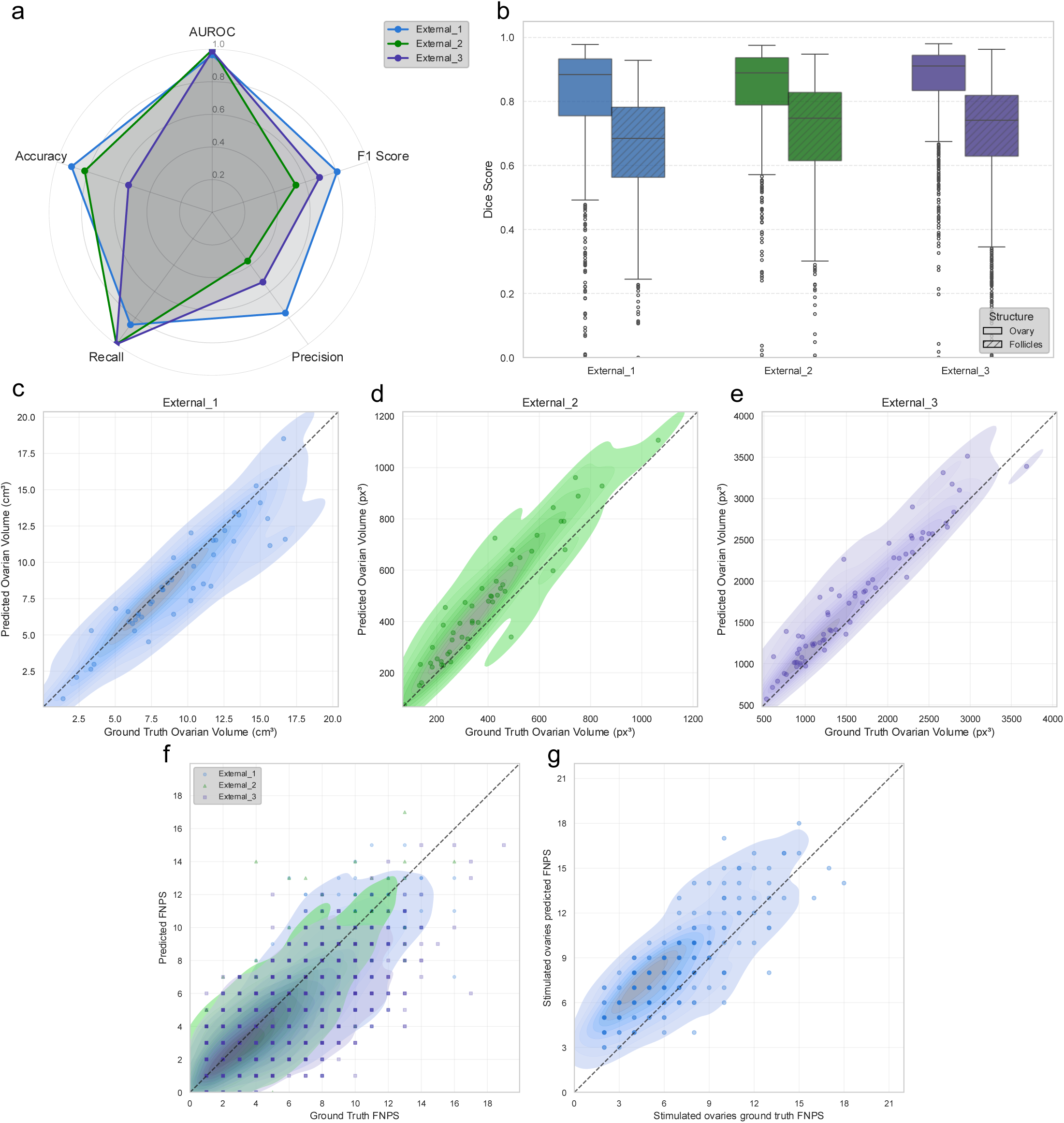
Performance of *Ovarify* on external validation datasets. (a) Frame classification performance on three external datasets (3D). (b) Ovary and follicle segmentation performance on three external datasets. (c-e) *Ovarify-*derived ovarian volume performance on External_1 (c), External_2 (d), and External_3 (e) datasets. Note that External_2 and External_3 datasets do not have pixel-cm conversion factors available to calculate volume in cm^3^. (f) *Ovarify-*derived FNPS performance on all frames in External_1, External_2, and External_3 datasets. (g) *Ovarify-*derived FNPS performance on External_4 dataset (single-frame stimulated ovaries). (c-g) Dashed lines represent perfect agreement.,

Finally, we applied the 2D *Ovarify* workflow (retrained on the entirety of the Cornell training dataset) to the *External_4* dataset, consisting of ovaries undergoing controlled ovarian stimulation garnered from clinical practice at the Center for Reproductive Medicine at Weill Cornell Medicine using the GE Voluson Signature 18 system. These single-frame ultrasonograms represent a distinct morphological regime characterized by dense, enlarged follicle populations absent from the training data and the prior external validation datasets. On ultrasonography, these follicles appear as large, tightly-packed anechoic structures, with thinner stromal borders separating follicles^34^. This morphology shift increases the difficulty of resolving individual follicular boundaries. Despite this change in morphological distribution, predicted FNPS correlated strongly with manual counts with a Pearson r of 0.813 (MAE 2.32 follicles/slice, n=268). This demonstrates that the *Ovarify*-learned representations capture follicular structure robustly enough to transfer to the stimulated-ovary setting encountered in assisted reproductive technology (Figure 4g).

## Discussion

Here, we developed and validated *Ovarify*, a novel deep learning workflow that comprehensively characterizes ovarian morphology directly from pelvic ultrasonography, producing ovarian and antral follicle segmentation masks alongside three clinically-established morphological measurements used to assess ovarian morphology: OV, FNPS, and FNPO^9, 11^. Across four-fold cross-validation, *Ovarify* segmented both the ovary and antral follicles with high fidelity, outperforming a baseline deep learning architecture. This segmentation performance translated into morphological assessments that closely agreed with manual assessment by trained annotators using validated image analysis protocols. We established that performance of *Ovarify* was preserved across four held-out datasets spanning distinct scanners, clinical settings, and populations. We additionally showed that *Ovarify* could be applied to a clinically-meaningful task of classifying polycystic ovarian morphology. Together, these results establish *Ovarify* as a reproducible, operator-independent tool for the quantitative assessment of ovarian morphology.

A significant barrier to the use of ovarian morphology has historically been the poor reproducibility of manual sonographic assessment, where inter-rater agreement for ovarian morphological metrics on pelvic ultrasonography is only moderate and follicle counting in the context of polycystic ovary morphology specifically is notably time-intensive and subjective^17, 19, 35–37^. Offline image analysis methods improve accuracy but demand specialized training and substantial time per scan, which has limited their clinical adoption^19–22^. *Ovarify* addresses both of these constraints simultaneously; by automating frame selection, segmentation, and follicle counting in a single end-to-end workflow, it eliminates the manual effort that has kept rigorous morphological analysis out of routine practice. This removes the operator-dependent steps that drive measurement variability. Additionally, because the classification stage uses a lightweight encoder and the workflow operates without manual frame selection, characterization of a full scan is achieved in a fraction of the time required for expert offline analysis, offering a path to standardized morphological assessment at a scale that manual methods cannot support. The ovary-to-follicle weight transfer further illustrates how anatomical structure can be exploited during training. This sequential design offers a simple strategy for hierarchical segmentation with minimal post-processing.

Notably, *Ovarify* is robust to input scan format. Adding three-dimensional context through adjacent-frame stacking achieved similar performance to single-frame input across all deep learning tasks and downstream measurements. This indicates that ovarian and follicular morphology can be characterized accurately from individual two-dimensional frames, and that the workflow can be applied out-of-the-box to static images and to frames sampled from manual cineloops or automated volumetric sweeps. This format-agnosticism is clinically significant, as it allows a single workflow to serve settings that acquire full 3D volumes as well as those that capture 2D images, including lower-resource environments where volumetric acquisition may be unavailable.

Ultimately, *Ovarify* characterizes ovarian morphology independent of ultrasonography hardware, clinical setting, and population. The workflow is trained and validated using reproducible image analysis techniques and demonstrates clinical utility in the context of PMOS-associated PCOM as well as ovarian stimulation. However, our study has several limitations. First, the primary training cohort was drawn from a limited number of contributing centers under research data collection protocols with specified image presets, and although external validation demonstrated generalizability of the workflow, prospective evaluation in clinical populations will be necessary to establish real-world performance. Second, follicle-level measurements, particularly FNPO, showed greater error than ovarian volume, reflecting the intrinsic difficulty of resolving numerous small, tightly-packed follicles. Finally, the stimulated ovary analysis was restricted to FNPS on single-frame images from a single center, and broader evaluation across stimulation protocols and morphological metrics associated with assisted reproduction is required.

Several directions follow from this work. Connecting *Ovarify*’s morphological outputs to clinical endpoints, such as PMOS diagnosis, biochemical markers or androgen excess, and reproductive or cardiometabolic outcomes, would test whether automated morphological feature extraction can serve as a quantitative biomarker of disease severity rather than a diagnostic feature alone. Additionally, we have previously shown that a predictive framework combining ovarian volume with anti-müllerian hormone and a limited set of self-reported and examination-derived features achieves high diagnostic accuracy for PMOS, but this relies on ovarian volume obtained through operator-dependent offline analysis^38^. Supplying this measurement automatically would remove that bottleneck and allow such workflows to scale. The demonstrated transfer to stimulated ovaries also suggests utility in assisted reproductive technology, where reproducible follicle counts and measurements are central to monitoring ovarian response. Lastly, implementation of *Ovarify* on ultrasonographic hardware incorporated with follicle measurements would enable clinically-relevant point-of-care morphological measurements in an objective manner.

## Methods

### Curated pelvic ultrasonography datasets

*Ovarify* was trained and validated on multiple datasets representing a variety of research and clinical settings. The primary dataset (*Cornell dataset*) included retrospectively-sampled participants who were prospectively recruited to any one of four parent protocols between 2009 and 2021 (NCT01859663, NCT01785719, NCT01927471, NCT01927432) at one of three centers: University of Rochester (IRB# STUDY00000436, STUDY00001531), Cornell University (IRB# 1207003154, 1303003665, 0908000633, 1108002383), or Weill Cornell Medicine (IRB# 1410015577). A total of 986 3D transvaginal ultrasound scans were used under a standardized image collection protocol using established presets at each site, collected from 284 participants, evenly split across healthy or polycystic ovaries. All participants were aged between 18-48 years at the time of scan collection and depending on the recruiting study protocol, potentially have multiple scans included in the dataset. All ultrasounds in the Cornell dataset were performing using GE Voluson S– or E-Series (6, 8, and 10) systems with curved-array transvaginal transducers (RIC6-12-D, RIC5-9A-RS, and RIC5-9W-RS). Images were annotated for ovaries and antral follicles using an established image analysis workflow by trained annotators. Briefly, ovary and follicle segmentation masks were generated using Fiji (ImageJ). An in-house Python script and ImageJ macro prompted annotators to select the first and last frame number containing ovarian content. All frames outside this range were converted to negative masks. Annotators were shown intermittent frames within the preselected range and were prompted to trace and fill all ovarian or follicular content within each frame using a grid-based annotation method (grid size of 0.3 cm^2^) ^19^. Annotators were blinded to participant characteristics (including PMOS/PCOM status). Within the *Cornell dataset*, 44 participants (yielding 130 scans) were held-out for model inference. To validate FNPO calculation by *Ovarify* output, a further subset of the held-out test dataset consisting of 34 participants (yielding 61 scans) was randomly selected to have all frames annotated.

In addition to the primary *Cornell dataset*, four external validation datasets (*External_1*, *External_2*, *External_3*, *External_4*) were used to assess *Ovarify*’s generalizability to sites, ultrasound model, and populations in the absence of further model training. *External_1* consists of 44 pelvic ultrasonograms garnered from 33 participants, each with polycystic ovaries, from the Kar Clinic & Hospital (Bhubaneswar, India). *External_2* consists of 70 pelvic ultrasonograms garnered from 53 participants visiting the University of Saskatchewan (Saskatoon, Canada), with manual cineloops of both healthy and polycystic ovaries. *External_3* consists of a clinical cohort of 54 ultrasounds garnered from 33 participants with both healthy and polycystic ovaries, presenting to the University of Rochester Medical Center with fertility concerns. *External_4* consists of 367 single-frame pelvic ultrasounds from patients undergoing ovarian stimulation at Weill Cornell Medicine-New York Presbyterian. The *External_2*, *External_3*, and *External_4* validation datasets were imaged using Ultrasonix Sonix RP, GE Voluson S-Series, and GE Voluson Signature 18 systems, respectively, all with transvaginal transducers. The *External_1* validation dataset was imaged using a mixture of GE Voluson E8 or E10-series systems with a transvaginal transducer (RIC6-12-D). With the exception of *External_4*, all scans in the external validation datasets were annotated similarly to the *Cornell dataset*; *External_4* was annotated specifically for antral follicles in all scans.

All scans used for this study were deidentified by the respective collecting institutions and shared under cross-institutional data use and sharing agreements. The retrospective use of deidentified ultrasonograms was considered exempt from IRB review (IRB0149648), in accordance with Cornell University IRB policy.

### Scan pre-processing

In order to maintain faithfulness to the original ultrasound scans, we performed minimal pre-processing. All ultrasonograms were deidentified by the respective institution, and A– and B-planes were extracted in DICOM format. For model input, we further extracted individual frames from each 3D ultrasonogram in either jpeg or tiff format and uniformly cropped each frame to remove any machine-specific and institution-specific text. For adjacent-frame input, adjacent frames (single-channel grayscale) are concatenated depth-wise to create a 3-channel image, where the middle channel is the frame of interest. For single-frame input, a single-channel grayscale image is triplicated to create a 3-channel image. All frames are resized to 288×288 pixels.

### *Ovarify* deep learning method

*Ovarify* is a deep learning method designed to characterize ovarian morphology in healthy and polycystic ovaries using three stages: (1) a preliminary classifier, (2) ovarian segmentation, and (3) antral follicle segmentation.

In the preliminary classifier, an EfficientNet-b2 encoder takes in a 3-channel input as either a static frame (single grayscale image triplicated across all channels) or three adjacent frames from a multi-frame ultrasound^39^. An output dimension of 1 provides a probability of a frame containing ovarian content.

Ovarian and follicle segmentation are performed using a U-Net-inspired segmentation architecture^40^. For all frames that are positively classified as having ovarian content, a Xception encoder and subsequent MA-Net decoder with an output dimension of one channel predicts a binary ovarian mask and in parallel, a flattened antral follicle mask^41, 42^. Predicted segmentation masks are used for the post-model calculation of ovarian volume and antral follicle count in both 2D and 3D.

### Morphological metrics

Ovarian volume was computed from the per-frame ovarian segmentation masks. For each frame, the segmented ovarian area was obtained as the sum of foreground pixels and converted to physical units using an ultrasound-specific conversion factor (CF, in mm per pixel), giving a slice area in mm². To account for frames in which no ovarian mask was produced, masks for missing frames between two segmented frames were reconstructed by linear interpolation, with each interpolated frame formed as a distance-weighted combination of the two flanking segmented masks. The full set of observed and interpolated slices was then integrated across the sweep, scaling the summed pixel volume by CF² and by the inter-slice spacing to yield ovarian volume in cm³. This approach treats the ultrasound sweep as a stack of parallel sections and estimates volume by summation across sections rather than by an ellipsoid approximation, avoiding the geometric assumptions that constrain the conventional prolate-ellipsoid formula^35, 43^.

Follicle counts were derived from the per-frame binary follicle segmentation masks. For each frame, a Euclidean distance transform was computed over the binary mask, and local maxima of the distance transform were identified as follicle centroids using a minimum separation of 10 pixels, with detection restricted to within segmented follicle regions. The number of centroids in a single frame defined the follicle number per single slice (FNPS). To compute the follicle number per ovary (FNPO), centroids from all frames were assembled into a three-dimensional volume in which each detected centroid was marked, preserving its frame index and in-plane position. Because a single follicle spans multiple contiguous frames and may shift slightly between slices, the centroid volume was dilated with an anisotropic structuring element that connected detections across ±4 pixels in plane and ±2 frames in depth. Connected-component labeling of the dilated volume, using full connectivity, then merged centroids belonging to the same physical follicle; FNPO is then defined as the resulting number of connected components. This three-dimensional reconciliation prevents the repeated counting of individual follicles across adjacent frames that would otherwise inflate per-ovary counts derived from independent two-dimensional slices.

### Training *Ovarify*

For each respective stage of *Ovarify*, we applied the same training recipe to both the single-frame as well as adjacent-frame model variants. The classification stage of *Ovarify* was trained for 100 epochs with a cosine annealing learning rate scheduler with warm restarts (learning rate of 1×10^−5^ with a 25-epoch period) and a binary cross-entropy loss objective. Both segmentation stages of *Ovarify* were trained using a Jaccard loss objective. Ovarian segmentation was trained for 180 epochs with a cosine annealing learning rate scheduler with warm restarts (learning rate of 1×10^−4^ with a 60-epoch period). Follicle segmentation was trained for 200 epochs with a custom learning rate scheduler; after a cosine warmup for 75 epochs to a learning rate of 5×10^−5^, the learning rate is held constant until epoch 175 when it follows an exponential decay with a decay rate of 0.8. Classification is optimized using stochastic gradient descent, and both segmentation stages using AdamW. Ovarian segmentation and follicle segmentation are trained sequentially, with follicle segmentation weights being initialized using converged ovarian segmentation weights. The classification and ovarian segmentation weights are trained with ImageNet-1k weight initializations. Across all stages of *Ovarify*, the following augmentations were applied to supplement the training dataset: horizontal/vertical flips, affine rotations, distortions, and noise and contrast augmentations.

### Statistics, validation, and reproducibility

Stages of *Ovarify* were architecturally validated against a baseline CNN. For the classification stage, *Ovarify* was validated against ResNet-34, pre-trained using ImageNet-1k, with an output dimension of 1^44^. For the segmentation stages, we constructed a U-Net segmentation model with a ResNet-34 backbone and followed a similar weight transfer paradigm in training the ovarian segmentation and follicle segmentation stages. To maintain a fair evaluation, the baseline workflow was trained to convergence using the same training recipe as *Ovarify*.

All model evaluations using the *Cornell dataset* evaluations were performed using four-fold cross-validation, with folds randomly partitioned at the patient level to ensure that all scans from a given participant appeared only in a single training fold. Differences in performance between model configurations and baselines were performed using the Wilcoxon signed-rank test apart from comparing classification AUROC; classification AUROC was evaluated for statistical significance using DeLong’s test. Statistical significance was defined as p<0.05. For comparison of classification metrics, significance was calculated with bootstrapping (n=1000). Classification metrics (except AUROC) are all reported at classification threshold of 0.5. All model development and analyses were performed using Python (v3.10.9), with model development using PyTorch (v1.13.1), and Albumentations (v1.3.1). Additionally, statistical analysis used SciPy (v1.10.1). All randomization seeds were fixed for reproducibility.

In generating ground truth segmentation masks, annotators were blinded to PCOM and PMOS status and assisted reproductive technology outcomes, in addition to all associated participant and scan metadata. Investigators were not blinded. Annotators received comprehensive training in the identification of the ovary and antral follicles on pelvic ultrasonography and were evaluated on their ability to calculate OV and FNPO from a subset of scans from the *Cornell dataset* using a grid-based method^19^. Reproducibility of annotators in analyzing pelvic ultrasounds was evaluated using Intraclass Correlation Coefficient (ICC).

### Resource requirements

All models in the *Ovarify* workflow were trained on 2x NVIDIA RTX A6000 GPUs, housed on a high-performance computing cluster at the BioHPC Bioinformatics Facility at Cornell University. On the full training dataset, the classification stage of *Ovarify* was trained to convergence in (hh:mm:ss) 53:22:15 (95% CI, 36:32:37-70:11:53) and used a peak of 13.0 GB of RAM. The ovarian and follicle segmentation stages were trained to convergence in 11:59:44 (95% CI, 11:24:47-12:34:40) and 9:57:13 (95% CI, 4:55:41-14:58:46), respectively; both models used a peak of 10.6 GB of RAM. The end-to-end inference workflow on one 3D ultrasonogram, using a single NVIDIA RTX A6000 GPU processing each frame serially, took a total time of 3.170 s (2.042 s [95% CI, 1.987-2.097 s], 0.545 s [95% CI, 0.524-0.566 s], and 0.583 s [95% CI, 0.562-0.605 s] for classification, ovarian segmentation, and follicle segmentation, respectively) using a peak of 3.4 GB of RAM (n=61 scans, fully-labeled FNPO subset of Cornell held-out test dataset), highlighting the low resource requirements of the *Ovarify* workflow.

## Author Contributions

ER, KH, ML, and IH conceived the study. ER, MB, and JP performed model prototyping. FC, BS, JP, SS, KH, and ML provided model training data and internal held-out test data. AE, JM, SK, DC, RP, SS, KH, and ML provided validation datasets. ER developed the model and performed all computational analyses and validations. AE, KH, and SS provided clinical data interpretation. ER wrote the manuscript and created figures, with input from FC, KH, ML, and IH. All authors provided input in editing the manuscript. IH and ML supervised the study, with input from KH. ER, JP, AE, KH, ML, and IH secured funding. ER, BS, and IH have personally accessed and verified the data. All authors have full access to all data in the study and accept responsibility to submit for publication.

## Acknowledgements

This work was supported by NICHD 1R21HD116139 (K.H., M.L., I.H.), Richard W. and Mae Stone Goode Foundation Research Award (A.E., K.H.), Cornell University CALS Moonshot Seed Grant (M.L., I.H.), Cornell University Multi-Investigator Seed Grant CUI 228215-01 (M.L., I.H.), and NCATS TL1TR002386 (E.R., J.P.). The funding played no role in study design, data collection, analysis and interpretation of data, or the writing of this manuscript. The authors additionally extend their gratitude to trained annotators in the Cornell Ovary Lab and at the University of Rochester Medical Center.

## Competing interests

IH is a consultant and on the advisory board of Noor Sciences. The other authors declare no competing interests.

## Data availability

Owing to reasonable privacy and security concerns, the primary (*Cornell*) training dataset and the external pelvic ultrasonography datasets (*External_1*, *External_2*, *External_3*, *External_4*) used in this study are not publicly available. We have provided the ability to request access to our held-out test dataset on Zenodo (https://doi.org/10.5281/zenodo.22259179).

## Code availability

The source code for the *Ovarify* deep learning workflow, in addition to morphological metric calculation and image annotation workflows, are available on GitHub (https://github.com/ih-lab/Ovarify). The model weights are additionally available in the same repository, for academic use only.

## Notes

https://doi.org/10.5281/zenodo.22259179

