## Supplementary Material for "*Ovarify*: a generalizable deep learning method for the automatic characterization of ovarian morphology on pelvic ultrasonography"

### Supplementary Figure 1

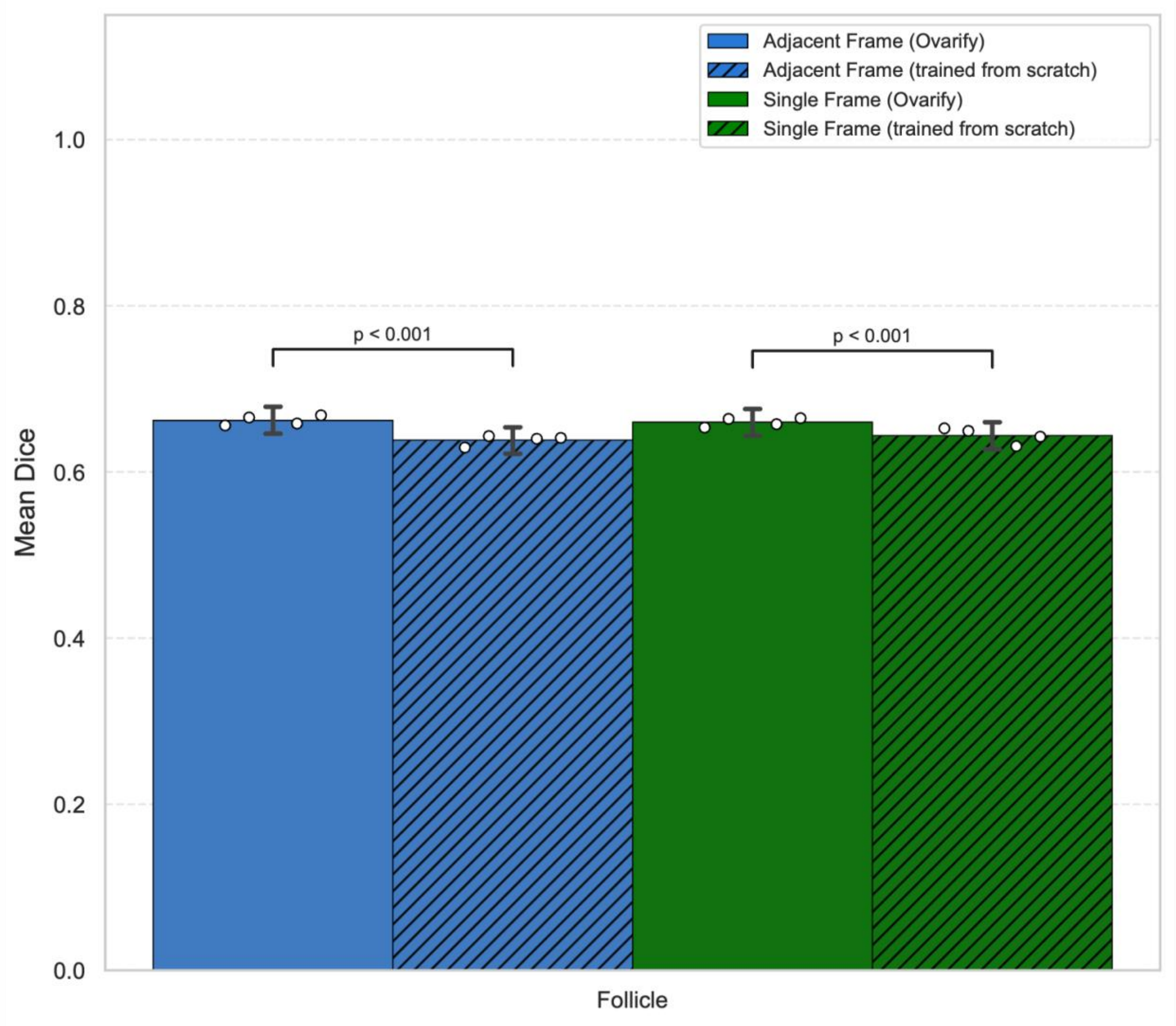


***Supplemental Figure 1***. Impact of training from scratch or training from *Ovarify* ovary segmentation converged weights for follicle segmentation. Error bars represent 95% confidence intervals across 4-fold cross-validation. Points represent mean Dice scores for individual folds.

### Supplementary Table 1

| Characteristic | Value |
|---|---|
| **Total participants** | 240 |
| **Age at ultrasound (years), n** | 237 |
| **Age at ultrasound (years), mean ± SD** | 27.9 ± 6.6 |
| **Age at ultrasound (years), median [IQR]** | 27.0 [22.0-33.0] |
| **Age at ultrasound (years), range** | 18.0-48.0 |
| **BMI (kg/m$^2$)**, n | 235 |
| **BMI (kg/m$^2$)**, mean ± SD | 28.7 ± 8.0 |
| **BMI (kg/m$^2$)**, median [IQR] | 26.7 [22.4-34.0] |
| **BMI (kg/m$^2$)**, range | 16.3-53.4 |
| **Race, n (%)** | |
| **White** | 154 (64.2%) |
| **Asian** | 34 (14.2%) |
| **Black or African American** | 26 (10.8%) |
| **Other or Unknown** | 17 (7.1%) |
| **More than one race** | 9 (3.8%) |
| **Ethnicity, n (%)** | |
| **Not Hispanic or Latino** | 193 (80.4%) |
| **Other or Unknown** | 28 (11.7%) |
| **Hispanic or Latino** | 19 (7.9%) |

***Supplementary Table 1.*** Demographic breakdown of participants in Cornell training dataset for *Ovarify*.

### Supplementary Table 2

| Characteristic | Value |
|---|---|
| **Total participants** | 44 |
| **Age at ultrasound (years), n** | 44 |
| **Age at ultrasound (years), mean ± SD** | 27.9 ± 5.5 |
| **Age at ultrasound (years), median [IQR]** | 27.0 [24.0-32.0] |
| **Age at ultrasound (years), range** | 19.0-40.0 |
| **BMI (kg/m$^2$)**, n | 43 |
| **BMI (kg/m$^2$)**, mean ± SD | 28.9 ± 9.3 |
| **BMI (kg/m$^2$)**, median [IQR] | 26.0 [21.9-35.3] |
| **BMI (kg/m$^2$)**, range | 15.7-51.7 |
| **Race, n (%)** | |
| **White** | 27 (61.4%) |
| **Black or African American** | 7 (15.9%) |
| **Asian** | 4 (9.1%) |
| **Other or Unknown** | 4 (9.1%) |
| **More than one race** | 2 (4.5%) |
| **Ethnicity, n (%)** | |
| **Not Hispanic or Latino** | 32 (72.7%) |
| **Hispanic or Latino** | 7 (15.9%) |
| **Other or Unknown** | 3 (11.4%) |

***Supplementary Table 2.*** Demographic breakdown of participants in Cornell held-out test dataset for *Ovarify*.

### Supplementary Table 3

| Structure | Measure | Intraclass Correlation Coefficient | 95% CI | n | k |
|---|---|---|---|---|---|
| Ovary | Single | 0.819 | [0.653, 0.941] | 10 | 9 |
| Ovary | Average | 0.976 | [0.944, 0.993] | 10 | 9 |
| Follicle | Single | 0.852 | [0.739, 0.937] | 15 | 9 |
| Follicle | Average | 0.981 | [0.962, 0.993] | 15 | 9 |

***Supplementary Table 3.*** Consistency analysis of annotators (k=9) in ovary (n=10) and follicle (n=15) identification after training in grid-based image analysis.

### Supplementary Table 4

| Model | Mean AUROC | AUROC 95% CI | Mean Precision | Precision 95% CI | Mean Recall | Recall 95% CI | Mean Accuracy | Accuracy 95% CI | Mean F1 Score | F1 Score 95% CI | N frames |
|---|---|---|---|---|---|---|---|---|---|---|---|
| **Adjacent Frame *Ovarify*** | 0.976 | [0.975, 0.977] | 0.616 | [0.551, 0.681] | 0.914 | [0.878, 0.950] | 0.903 | [0.882, 0.924] | 0.735 | [0.700, 0.770] | 10917 |
| **Single Frame *Ovarify*** | 0.969 | [0.963, 0.975] | 0.642 | [0.582, 0.702] | 0.887 | [0.833, 0.941] | 0.91 | [0.895, 0.925] | 0.743 | [0.719, 0.767] | 10917 |
| **Adjacent Frame ResNet-34** | 0.969 | [0.967, 0.971] | 0.629 | [0.563, 0.695] | 0.875 | [0.843, 0.907] | 0.905 | [0.885, 0.925] | 0.731 | [0.697, 0.765] | 10917 |
| **Single Frame ResNet-34** | 0.965 | [0.962, 0.968] | 0.587 | [0.525, 0.649] | 0.9 | [0.856, 0.944] | 0.891 | [0.870, 0.912] | 0.709 | [0.678, 0.740] | 10917 |

***Supplementary Table 4.*** Complete classification metrics for frame classification compared to baseline ResNet-34 across 4-fold cross-validation.

**Supplementary Table 5**

| Structure | Model | N frames | Mean Dice | 95% CI |
|---|---|---|---|---|
| Ovary | Adjacent Frame (*Ovarify*) | 1600 | 0.824 | [0.815, 0.833] |
| Ovary | Adjacent Frame (U-Net) | 1600 | 0.801 | [0.791, 0.811] |
| Ovary | Single Frame (*Ovarify*) | 1600 | 0.818 | [0.809, 0.827] |
| Ovary | Single Frame (U-Net) | 1600 | 0.793 | [0.783, 0.803] |
| Follicle | Adjacent Frame (*Ovarify*) | 741 | 0.662 | [0.646, 0.678] |
| Follicle | Adjacent Frame (U-Net) | 741 | 0.47 | [0.458, 0.483] |
| Follicle | Single Frame (*Ovarify*) | 741 | 0.66 | [0.644, 0.676] |
| Follicle | Single Frame (U-Net) | 741 | 0.506 | [0.493, 0.519] |

***Supplementary Table 5.*** Segmentation Dice scores for ovarian and follicle segmentation compared to baseline U-Net across 4-fold cross-validation.

**Supplementary Table 6**

| Model | N frames | Mean Dice | 95% CI |
|---|---|---|---|
| Adjacent Frame (*Ovarify*) | 741 | 0.662 | [0.646, 0.678] |
| Adjacent Frame (trained from scratch) | 741 | 0.638 | [0.623, 0.654] |
| Single Frame (*Ovarify*) | 741 | 0.66 | [0.644, 0.676] |
| Single Frame (trained from scratch) | 741 | 0.644 | [0.628, 0.660] |

***Supplementary Table 6.*** Segmentation Dice scores for follicle segmentation compared to random weight initialization for training across 4-fold cross-validation.

**Supplementary Table 7**

| Site | Metric | Mean | N frames | 95% CI |
|---|---|---|---|---|
| **External_1** | AUROC | 0.965 | 3519 | [0.958, 0.970] |
| **External_1** | F1 Score | 0.805 | 3519 | [0.786, 0.825] |
| **External_1** | Precision | 0.763 | 3519 | [0.739, 0.790] |
| **External_1** | Recall | 0.852 | 3519 | [0.827, 0.876] |
| **External_1** | Accuracy | 0.906 | 3519 | [0.897, 0.915] |
| **External_2** | AUROC | 0.995 | 7258 | [0.994, 0.997] |
| **External_2** | F1 Score | 0.539 | 7258 | [0.515, 0.561] |
| **External_2** | Precision | 0.369 | 7258 | [0.347, 0.390] |
| **External_2** | Recall | 0.999 | 7258 | [0.996, 1.000] |
| **External_2** | Accuracy | 0.821 | 7258 | [0.812, 0.830] |
| **External_3** | AUROC | 0.988 | 3150 | [0.984, 0.991] |
| **External_3** | F1 Score | 0.691 | 3150 | [0.676, 0.707] |
| **External_3** | Precision | 0.529 | 3150 | [0.510, 0.546] |
| **External_3** | Recall | 0.999 | 3150 | [0.998, 1.000] |
| **External_3** | Accuracy | 0.539 | 3150 | [0.521, 0.557] |

***Supplementary Table 7***. Complete frame classification metrics for three external validation datasets using *Ovarify* trained on entire training set.

**Supplementary Table 8**

| Dataset | Structure | N frames | Mean Dice | 95% CI |
|---|---|---|---|---|
| **External_1** | Ovary | 802 | 0.796 | [0.781, 0.811] |
| **External_1** | Follicles | 671 | 0.653 | [0.639, 0.666] |
| **External_2** | Ovary | 760 | 0.817 | [0.803, 0.832] |
| **External_2** | Follicles | 630 | 0.686 | [0.671, 0.702] |
| **External_3** | Ovary | 1629 | 0.867 | [0.860, 0.873] |
| **External_3** | Follicles | 2000 | 0.69 | [0.681, 0.698] |

***Supplementary Table 8.*** Segmentation Dice scores for ovary and follicle segmentation across three external validation datasets using *Ovarify* trained on the entire training set.

**Supplementary Table 9**

| Site | Pearson r | Error Metric | Error Value | Unit | N ovaries |
|---|---|---|---|---|---|
| **External_1** | 0.928 | MAE | 1.148 | $cm^2$ | 44 |
| **External_2** | 0.947 | MAPE | 24.315 | % | 53 |
| **External_3** | 0.960 | MAPE | 15.494 | % | 70 |

***Supplementary Table 9.*** Ovarian volume calculation performance of *Ovarify* on three external validation datasets.

**Supplementary Table 10**

| Site | Pearson r | MAE | N frames |
|---|---|---|---|
| **External_1** | 0.824 | 1.465 | 671 |
| **External_2** | 0.803 | 1.130 | 630 |
| **External_3** | 0.729 | 1.563 | 2000 |

***Supplementary Table 10.*** FNPS calculation performance of *Ovarify* on three external validation datasets.
